# Representational content of oscillatory brain activity during object recognition

**DOI:** 10.1101/2020.09.02.279216

**Authors:** Leila Reddy, Radoslaw Martin Cichy, Rufin VanRullen

## Abstract

Numerous theories propose a key role for brain oscillations in visual perception. Most of these theories postulate that sensory information is encoded in specific oscillatory components (e.g., power or phase) of specific frequency bands. These theories are often tested with whole-brain recording methods of low spatial resolution (EEG or MEG), or depth recordings that provide a local, incomplete view of the brain. Opportunities to bridge the gap between local neural populations and whole-brain signals are rare. Here, using representational similarity analysis we ask which MEG oscillatory components (power and phase, across various frequency bands) correspond to low or high-level visual object representations, using brain representations from fMRI, or layer-wise representations in Deep Neural Networks (DNNs) as a template for low/high-level object representations. The results showed that around stimulus onset and offset, most transient oscillatory signals correlated with low-level brain patterns (V1). During stimulus presentation, sustained beta (∼20Hz) and gamma (>60Hz) power best correlated with V1, while oscillatory phase components correlated with IT representations. Surprisingly, this pattern of results did not always correspond to low- or high-level DNN layer activity. In particular, sustained beta-band oscillatory power reflected high-level DNN layers, suggestive of a feed-back component. These results begin to bridge the gap between whole-brain oscillatory signals and object representations supported by local neuronal activations.

## Introduction

Oscillatory neuronal activity is thought to underlie a variety of perceptual functions. Different frequency bands can carry information about different stimulus properties (e.g., whether the stimulus consists of coarse or fine object features) (Smith, Gosselin et al. 2006, Romei, Driver et al. 2011), feedforward or feedback signals (van Kerkoerle, Self et al. 2014, Bastos, Vezoli et al. 2015), or may reflect neuronal communication between different neuronal populations (Fries 2005, Jensen and Mazaheri 2010). Other studies have shown that different components of an oscillation (e.g., its power or phase) encode different types of sensory information (Smith, Gosselin et al. 2006).

Although neuronal oscillations are observed in different brain regions, and key theories hold that they reflect processing within, and communication between, brain regions (Fries 2005, Jensen and Mazaheri 2010), it has been difficult to pin down how large-scale brain oscillations are related to local patterns of neural activity, and how this relationship unfolds over time. This is because oscillatory activity is often studied with methods such as EEG or MEG, which have low spatial resolution. Although oscillatory signals with high spatial specificity can be recorded via local field potential recordings in humans or animals, these methods usually only target specific brain regions, and thus can only provide a partial view of oscillatory activity and its role in large-scale brain function. A direct link between large-scale oscillations and local neural activity is missing.

Here, we combine large-scale oscillatory signals recorded by MEG with local patterns of neural activity recorded with fMRI to bridge the gap between oscillatory components and the different levels of object representation in the brain. Using representational similarity analysis (RSA, (Kriegeskorte, Mur et al. 2008)), we investigate the information carried by whole-brain oscillations obtained from MEG, and examine how this information evolves over time during an object recognition task.

We define three distinct dimensions of interest along which neural representations may unfold, and which are often conflated in the literature. First, we use the terms “early” and “late” to denote the *temporal* evolution of representations. Second, we differentiate between “low-level” and “high-level” stages of a processing *hierarchy*. Third, we consider the *complexity* of representations by distinguishing between “basic” and more “refined” information. In many information processing systems and in many typical experimental situations, these three dimensions are directly related to one another, as input information propagates over time through a succession of hierarchical stages, becoming more and more refined along the way. In such situations, the three dimensions of interest are in fact redundant and need not be further distinguished. But in systems with recurrence and feedback loops (like the brain), time, space and information complexity are not always linearly related. For example, a lower hierarchical level (e.g. V1) can carry more refined representations, later in time, as a result of feedback loops or lateral connections (Lamme and Roelfsema 2000). In our terminology, such a representation would be classified as late in time, low-level in the hierarchy, yet refined in terms of complexity.

In this work, we consider two main hierarchical systems. We are interested in understanding information processing in the human brain, so we use V1 and IT fMRI brain representations, as done in a number of recent studies (Cichy, Pantazis et al. 2014, Khaligh-Razavi and Kriegeskorte 2014). Representational similarity between MEG oscillations and this fMRI-based hierarchy can be interpreted in terms of early and late representations (based on the timing of the MEG oscillations), and in terms of low-level (V1) vs. high-level (IT) hierarchical stages. To assess the complexity of representations independent of temporal evolution and hierarchy of processing, we related our data to a second class of hierarchical systems: artificial feed-forward Deep Neural Networks (DNNs). In these artificial networks, the hierarchical level (low-level vs. high-level) is directly related to feature complexity (basic vs. refined representations), due to the absence of feed-back or recurrent loops. For any MEG oscillatory signal, representational similarity with DNN activation patterns can thus inform us about representational complexity. In turn, any difference between DNN-based and brain-based RSA may be suggestive of feed-back or recurrent influences in the MEG oscillatory signals.

With this dual approach, we find a complex picture of transient and sustained oscillatory signals that can be related to V1 and IT representations. Transient oscillatory components around stimulus onset and offset, as well as sustained beta (∼20Hz) and gamma (>60Hz) power components resemble V1 representations, while phase-dependent sustained activity correlates best with IT representations. However, when compared to DNNs, some early V1-like components actually correlate more with higher DNN layers, suggesting that stimulus representations early in time may already include refined information, presumably as a result of feedback or top-down influences (Kar, Kubilius et al. 2019, Kietzmann, Spoerer et al. 2019).

In effect our results narrow the gap between the description of neural dynamics in terms of whole-brain oscillatory signals and local neural activation patterns. Disentangling temporal evolution, hierarchical stage of processing and complexity of representations from each other, our approach allows for a more nuanced view on cortical information flow in human object processing.

## Methods

### Experimental paradigm and data acquisition

The data analyzed in this study was obtained from (Cichy, Pantazis et al. 2014), and detailed methods can be obtained from that paper.

Fifteen subjects performed separate MEG and fMRI sessions while they viewed a set of 92 images. The image set consisted of human and non-human faces and bodies, and artificial and natural everyday objects. The 92-image stimulus set was taken from the Kiani image set (Kiani, Esteky et al. 2007), which consists of cutout objects on a gray background.

In the MEG sessions, each image was presented for 0.5s followed by an inter-stimulus interval (ISI) of 1.2 or 1.5s. Every 3-5 trials, a target paperclip object was presented, and subjects’ task was to press a button and blink whenever they detected this target image. Subjects performed 2 MEG sessions, of 2 hours each. In each session they performed between 10 to 15 runs. Each image was presented twice in each run, in random order.

In each of two fMRI sessions, each image was presented for 0.5s followed by an ISI of 2.5 or 5.5s. Subjects’ task in the fMRI sessions was to press a button when they detected a color change in the fixation cross on 30 null trials, when no image was presented. Each image was presented once in each fMRI run, and subjects performed 10-14 runs in each session.

The MEG data was acquired from 306 channels (204 planar gradiometers, 102 magnetometers, Elekta Neuromag TRIUX, Elekta, Stockholm) at the Massachusetts Institute of Technology. The MRI experiment was conducted on a 3T Trio scanner (Siemens, Erlangen, Germany), with a 32-channel head coil. The structural images were acquired using a T1-weighted sequence (192 sagittal slices, FOV = 256mm^2^, TR=1,900ms, TE=2.52ms, flip angle=9 degrees). For the fMRI runs, 192 images were acquired for each participant (gradient-echo EPI sequence: TR = 2,000ms, TE=32 ms, flip angle = 80 degrees, FOV read = 192 mm, FOV phase = 100%, ascending acquisition gap = 10%, resolution = 2mm, slices=25).

### MEG analysis - preprocessing

MEG trials were extracted with a 600 ms baseline before stimulus onset until 1200 ms post-stimulus onset. A total of 20-30 trials were obtained for each condition, session, and participant. Each image was considered as a different condition.

Data were analyzed using custom scripts in Matlab (Mathworks) and FieldTrip (Oostenveld, Fries et al. 2011). Data were downsampled offline to 500 Hz. For each trial and sensor, we computed the complex time frequency decomposition using multitapers. Parameters used were: 50 distinct frequencies increasing logarithmically from 3 to 100 Hz, over a time interval of −600ms to 700ms with respect to stimulus onset, in steps of 20 ms. The length of the sliding time window was chosen such that there were two full cycles per time-window. The amount of smoothing increased with frequency (0.4 * frequency).

From the complex number at each time-frequency (TF) coordinate, we extracted two measures for each sensor and each condition: the power and the phase of the oscillation. For each channel and condition, on each trial, the power was first expressed in decibels, and then averaged across trials to obtain one power value per condition. The phase of the oscillation was obtained by first normalizing each trial to make each trial’s vector in the complex domain of unit length, and then averaging across trials for each condition. The resultant average vector was then normalized to unit length, and the sine (real) and cosine (imaginary) components were extracted for each condition and each sensor.

### MEG analysis – multivariate analysis (Figure 1a)

At each TF coordinate and for each condition, we next arranged the 306 power values from the 306 MEG sensors into a 306-dimensional vector representing the power pattern vector for that condition. Similarly, at each TF coordinate and for each condition we concatenated the 306 sine and 306 cosine values into a 612-dimensional phase pattern vector for that condition.

**Figure 1:**
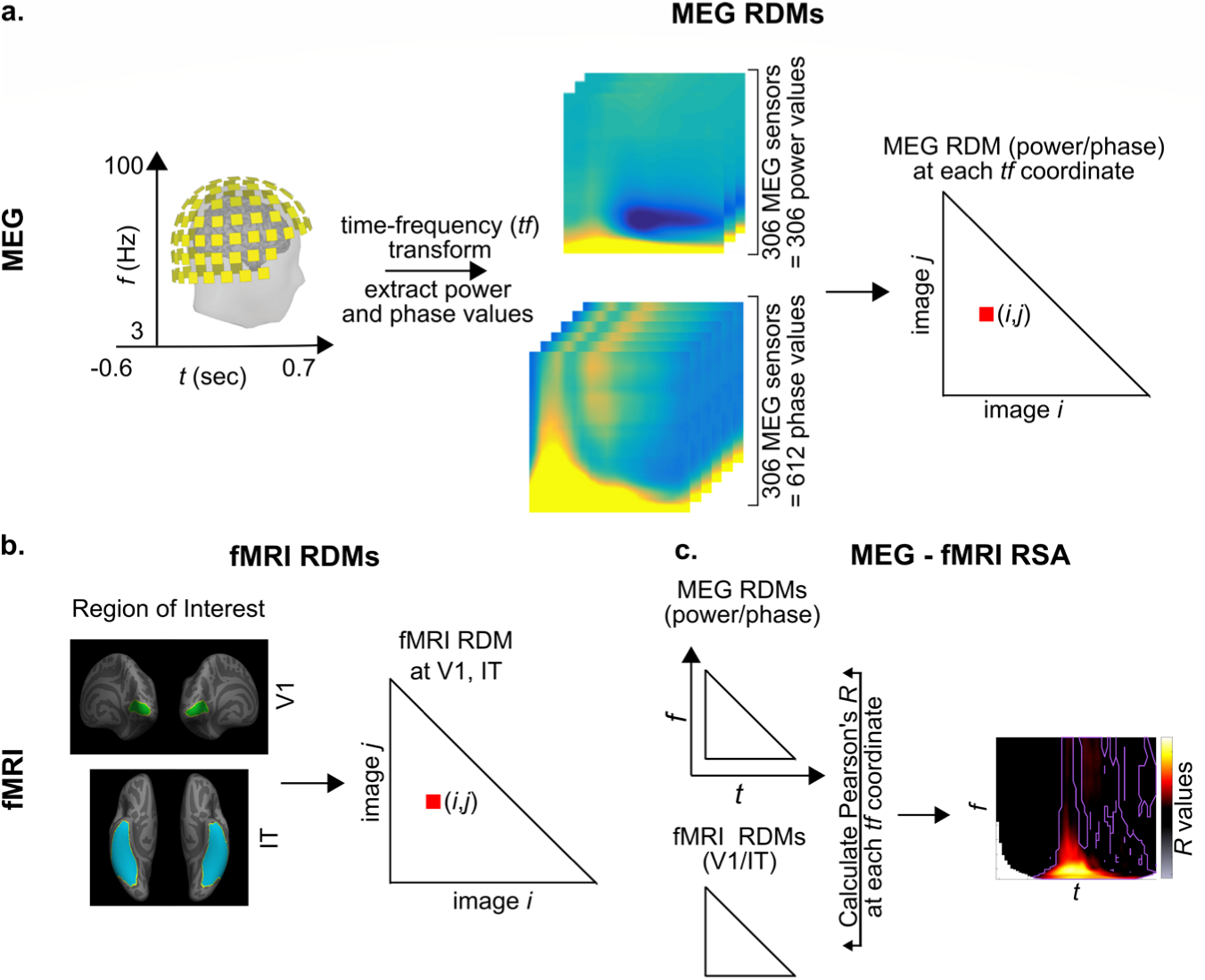
MEG-fMRI RSA analysis. (a) MEG analysis and MEG representational dissimilarity matrices (RDMs). From the MEG signals, the complex time frequency (TF) transform was computed for each of the 306 MEG sensors. The amplitude and phase (separated into cosine and sine) values were extracted from the complex number at each TF coordinate, and a MEG RDM was constructed, reflecting the distance between oscillatory activation patterns for every pair of images (*i,j*) (see methods for details). As a result, we obtained a power and phase MEG RDM at each TF coordinate for each participant. (b) fMRI RDMs were obtained from (Cichy, Pantazis et al. 2014). Two regions of interest (ROI) were defined: V1 and IT and one fMRI RDM was obtained for each ROI, and each participant, reflecting the distance between BOLD activation patterns for every pair of images (*i,j*). (c) The MEG power or phase RDMs were compared to the fMRI RDMs (V1 or IT) by computing the partial Pearson’s *R*. This step was performed at each TF coordinate, resulting in an RSA map of R values at each TF coordinate, for each subject and ROI.

We next computed two representational dissimilarity matrices (RDMs): one for power and one for phase, at each TF point. For each pair of conditions, the power (phase) pattern vectors were correlated using the Pearson correlation measure, and the resulting 1-correlation value was assigned to a 92 by 92 power (phase) RDM, in which the rows and columns corresponded to the images being compared. This matrix is symmetric across the diagonal. This procedure results in one power (phase) RDM at each TF point.

### fMRI analysis (Figure 1b)

The preprocessing steps for the fMRI data are described in detail in (Cichy, Pantazis et al. 2014). For the multivariate analysis, two regions of interest (ROIs) were defined: V1 and IT. In each subject, for each ROI, voxel activation values were extracted for each condition, and the resulting values were arranged in a pattern vector for each condition. Then, in each ROI, for each pair of conditions, the corresponding pattern vectors were correlated using the Pearson correlation measure, and the resulting 1-correlation value was assigned to the 92×92 fMRI RDM. For further analysis the fMRI RDMs were averaged across the 15 subjects, resulting in one RDM per ROI. The fMRI RDMs were provided by R. Cichy, D. Pantazis and A. Oliva (Cichy, Pantazis et al. 2014).

### MEG-fMRI Representational Similarity Analysis (RSA) (Figure 1c)

RSA between the MEG and fMRI RDMs was performed by computing the partial Pearson’s correlation between each MEG (phase or power) RDM with each fMRI RDM (V1 or IT), while partialling out any contribution from the other fMRI RDM (IT or V1). We chose to perform a partial correlation because the V1 and IT RDMs were positively correlated with each other (*r*∼0.3); compared to a standard correlation, the partial correlation allowed us to isolate the unique correlation of each fMRI RDM with the MEG RDM, while discarding their joint contribution.

This procedure resulted in four RSA maps (power/phase MEG RDMs x V1/IT fMRI RDMs). Each RSA map shows the R-value between the MEG signals and the V1/IT activation patterns at each TF point. Significance of the RSA result was evaluated with a paired t-test against 0, FDR corrected, alpha = 0.05.

### Clustering analysis

The MEG time-frequency RDMs are heavily correlated with each other. To facilitate the interpretation of the information content of oscillatory signals, and to determine which features co-vary and which are independent, K-means clustering was performed on the MEG power and phase RDMs. Clustering was performed on the 4186-dimensional ((92*92-92)/2) RDMs across all (66) time points and (46) frequency points, combining the power and phase signals (resulting in 46*66*2 = 6072 data points to cluster in a 4186-dimensional space). K-means was implemented with the Matlab function kmeans, with the correlational distance measure, five replicates, and the number of clusters going from 1 to 20. The optimal number of clusters was then determined with the elbow criterion defined as the point just before the local maximum of the second derivative of the residual sum of squares (corresponding to the point at which adding another cluster would only provide a marginal gain in variance explained). With this method, the first elbow, occurred at *k*=7 clusters.

The chosen clusters could be visualized by plotting the correlation distance (in the 4186-dimensional RDM space) between the cluster’s centroid and every time-frequency point, for both power and phase signals.

RSA (using partial Pearson’s correlation) was performed between each cluster’s centroid and each of the fMRI RDMs (see below). For RSA with fMRI, this procedure resulted in two RSA values (one each for V1 and IT). Since each cluster centroid could correspond to both V1 and IT to different degrees, the information content of the cluster was positioned somewhere between V1/low-level and IT/high-level using the following equation:

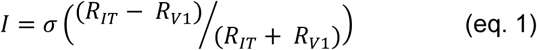

where σ denotes the sigmoid function. The measure *I* could vary between 0 (when the cluster’s representational content was perfectly similar to V1) and 1 (when it was perfectly similar to IT).

Significance of RSA between the cluster centroids and the fMRI RDMs was computed with a surrogate test. On each iteration, the cluster centroid RDM was randomly shuffled and the partial correlation was computed between this shuffled RDM and the true RDM. This procedure was repeated for 10^5^ iterations, and the number of iterations on which the shuffled RSA values were higher than the true RSA values was counted.

### Deep Neural Network (DNN) RDMs

The MEG phase/power representations were also compared to representations in four DNNs (so as to ensure that conclusions were not dependent on one specific network architecture): AlexNet (Krizhevksy, Sutskever et al. 2012), VGG16 (Simonyan and Zisserman 2014), GoogleNet (Szegedy, Liu et al. 2015), and InceptionV3 (Szegedy, Ioffe et al. 2017) processing the same 92 images as in our MEG and fMRI data. However, in contrast to our 92-image stimulus set, which consisted of cutout objects on a gray background, the DNNs had been trained on images from ImageNet (millions of photographs with one or more objects in natural backgrounds). The networks had thus learned optimal representations for their training set, but in this representation space our 92 images tended to cluster into a remote “corner” (Figure 2), with low dissimilarity (1-Pearson’s *R*) values between images, and a resulting RDM of poor quality. To retrieve meaningful distances between the representations of the 92 images, we first performed a centering procedure: we centered the activation of each layer of each DNN by subtracting the mean activation of an independent set of 368 images from the Kiani image set. This independent image set consisted of four images from each of the categories in our 92-image set. Importantly, because the image set used for centering did not include any of the 92 images from our study, there was no circularity in the centering operation, nor any leakage of information between the representations of our 92 images.

**Figure 2:**
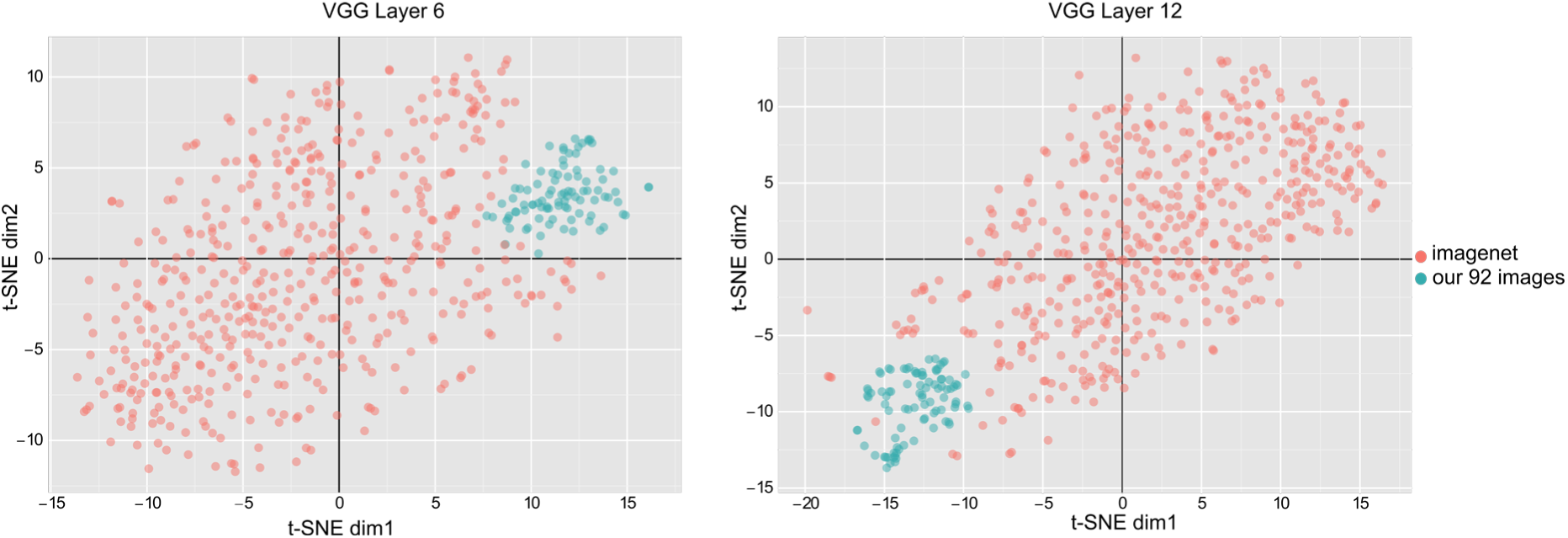
t-SNE visualizations of 500 ImageNet samples and the 92-image stimulus-set used in this study across two representative layers of the VGG network. The DNNs used in this study had been trained on images from ImageNet, which consists of millions of photographs of one or more objects in natural backgrounds. In contrast, our 92-image stimulus set consists of cut-out images on a gray background. The DNNs learn optimal representations for the training images from ImageNet, i.e., different images from different categories are mapped to different regions of the representation space, and the whole space tends to be equally occupied by the training samples. However, as the t-SNE visualizations show, our 92 images are all projected into a remote corner of this space, meaning that the RDM distances between the 92 images are confounded by the mean vector (the pairwise Pearson distance depends more on the alignment with the mean vector, and less on the true physical distance between points). To circumvent this problem, we used a re-centering approach as described in the methods section.

RDMs were constructed for each convolutional layer of each network based on the layer activation values. There were 5 layers for AlexNet, 13 for VGG16, 12 for GoogleNet and 16 for InceptionV3. RSA was then performed (with the Spearman correlation) between these RDMs and the centroid of each cluster (see above for details of the clustering analysis). The layer with maximum RSA, normalized by the number of layers in the DNN, was taken to reflect the information content of this cluster between 0 (in the terminology defined in the Introduction, “low-level” and basic, corresponding to the DNN’s first layer) and 1 (“high-level” and refined, corresponding to the DNN’s last layer), and finally averaged across the four DNNs.

## Results

Fifteen participants viewed the same set of 92 images while fMRI and MEG data was recorded (in separate sessions). The image set consisted of human and non-human bodies and faces, and artificial and natural stimuli. Each stimulus was presented for 0.5s, followed by a 1.2 or 1.5s baseline period.

To assess oscillatory components, we extracted stimulus-related activity from −600ms to 1200ms relative to stimulus onset from the MEG data. For each trial, and each sensor a time-frequency (TF) decomposition was performed, and a power and phase value extracted at each time and frequency point. These values were used to compute representational dissimilarity matrices (RDMs) at each TF point, separately for power and phase (Methods and Figure 1a). Each element in the MEG RDMs indicates how distinct the corresponding images are in the MEG power or phase spaces, and the entire MEG RDM is a summary of how the 92-image stimulus set is represented in the MEG oscillatory power or phase at each TF point.

To assess local patterns of neural activity we generated fMRI RDMs by performing comparisons between the local BOLD activation patterns of pairs of images in V1 and IT (Cichy, Pantazis et al. 2014). Two fMRI RDMs were obtained (Figure 1b), one for V1 and one for IT. The fMRI RDMs are a measure of the representation of the image set in the voxel space of V1 and IT local neural activity.

### Bridging the space, time and frequency gap in object recognition

How similar is the oscillatory representation of the images to their representation in each brain region? The MEG RDMs (power and/or phase) at each TF point represent the stimulus set in a large-scale brain oscillatory activity space, while the fMRI RDMs represent the same image set via BOLD activity in a local population of neurons in two brain regions (V1 or IT). We evaluated the similarity of representations in the time-frequency domain with those in the fMRI activation patterns by computing the partial Pearson’s correlation between the MEG RDMs (phase or power) with the fMRI RDMs (V1/IT), at each TF point (Figure 1c). This analysis resulted in four time frequency maps of R-values (or RSA maps), which provide the unique correspondence between whole-brain oscillations and local patterns of neural activity in V1 and IT, at each TF point (Figure 3). With these maps we can ask if and when stimulus information contained in oscillatory phase or power at each TF point resembles BOLD activations in a given brain region (V1/IT), and potentially, which region it resembles more. The advantage of measuring partial correlation (instead of a standard correlation) is to discard the (potentially large) portion of the variance in oscillatory representations that is explained equally well by V1 or IT BOLD representations— owing to the fact that V1 and IT signals already share similarities. This way, we concentrate on the part of oscillatory representations that is *uniquely* explained by each brain region-of-interest.

**Figure 3:**
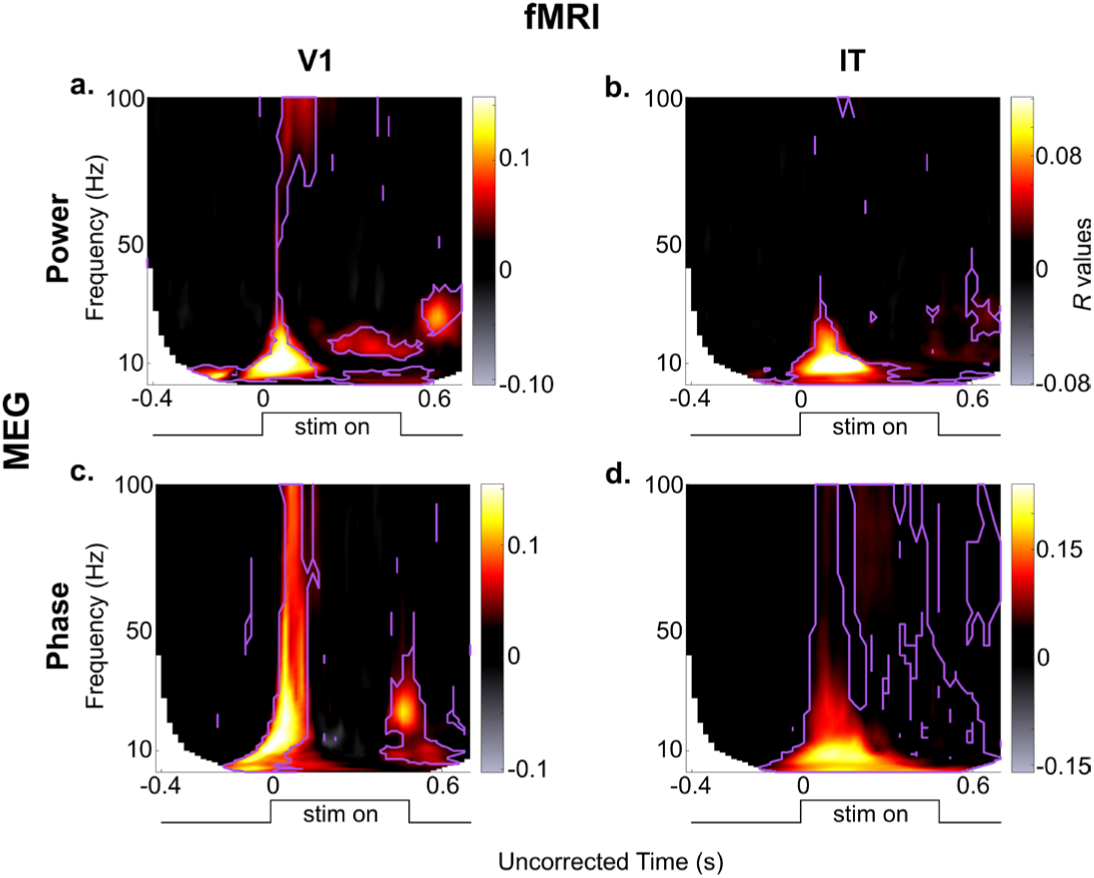
Results of the 2×2 RSA comparisons (MEG power/phase x fMRI V1/IT), averaged over all subjects. The purple contours mark those regions in the maps that are significantly different from zero (paired *t-* test against 0 across *N*=15 subjects, FDR correction, alpha = 0.05). Note that the absolute latencies are not directly comparable across frequencies, because of different smoothing windows applied at the different frequencies when performing the TF transform (hence, the x-axis is labeled as uncorrected time).

Our results show that different oscillatory components map to different brain regions at different moments in time. Overall, the absolute maximum of representational similarity with brain area V1 occurred in the alpha band around 120ms post-stimulus for oscillatory power, whereas the absolute maximum related to area IT occurred for theta- and alpha-phase around 200-300ms. More generally, a strong increase in representational similarity was observed shortly after stimulus onset in all four maps. The frequency, latency and duration of these similarity effects depended however on the exact oscillatory signal (power, phase) and brain region (V1, IT). In terms of MEG power (Figure 3a, b), the latencies (see also Figure 4) respected the hierarchical order of visual processing (Nowak and Bullier 1998) with an increase in representational similarity in the lower (<20 Hz) frequency bands occurring around the evoked response first for V1, and about 20-30ms later for IT (paired t-test against 0, FDR corrected, alpha=0.05). This latency difference is similar to that reported in (Cichy, Pantazis et al. 2014), where the peak correspondence between the average MEG signal and V1 activity occurs about 30 ms prior to the peak with IT activity. The onset response in V1 also consisted of high gamma frequencies (>70Hz), whereas this high-gamma activity was not observed in IT. A sustained low-beta (20Hz, 200-500ms) and an offset high-beta (30Hz, ∼600ms) response also corresponded to V1 representations, although neither of these effects were observed in IT (see also Figure 4). In terms of stimulus representations in the MEG oscillatory phase (Figure 3c, d), after an initial broadband (3-100Hz) transient peak at stimulus onset corresponding to V1 representations, stimulus information carried by sustained oscillatory phase resembled IT representations in the low (<20Hz) and high frequency (60Hz) bands, and this resemblance persisted until the end of the trial. Phase representations corresponding to V1 patterns were observed again around stimulus offset, at alpha (∼10Hz) and beta (20-30Hz) frequencies (see also Figure 4).

**Figure 4.**
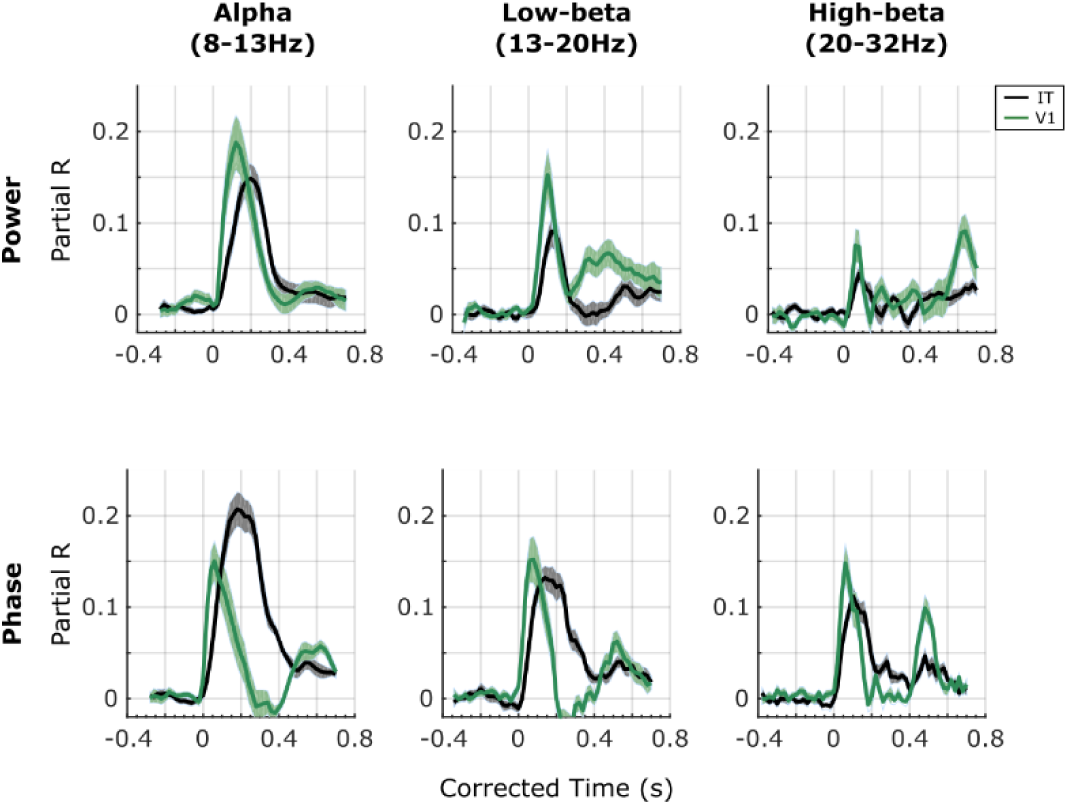
Profile of the results of the RSA with V1 (green lines) and IT (black lines) in oscillatory power (top row) and oscillatory phase (bottom row) in different frequency bands. To examine the RSA maps in more detail, we extracted their time courses in different traditional frequency bands: alpha (8-13 Hz), low-beta (13-20 Hz), and high-beta (20-32 Hz). In each of these frequency bands we computed the average R-values. Since the TF decomposition induces temporal smearing, and the amount of smearing differs for different frequencies, in order to interpret the latencies of the representational similarities, we corrected for this smearing effect. Specifically, to avoid underestimating the onset latencies, we corrected time by adding half the wavelet window duration at each frequency. Note that the same correction was applied to the two curves compared in each plot. Solid lines are the means across subjects, and the shaded areas correspond to the SEM across subjects.

These results thus suggest that different oscillatory components correspond to different brain regions at different time-frequency points. However, since the RDMs in the time-frequency space are heavily correlated with each other, it is difficult to ascertain from this analysis which power/phase features co-vary, and which effects occur independently. To better interpret the results shown in Figure 3 we turned to a clustering analysis. The clustering analysis allowed us to reduce the dimensionality of the dataspace and to determine which oscillatory signals occurred jointly, and which are independent. We performed k-means clustering jointly on the power and phase RDMs. The results of the clustering analysis for *k*=7 clusters (the “optimal” number of clusters for our dataset) are shown in Figure 5 (see also Figure 6). The first cluster (ranked by smallest distance from cluster centroid) corresponded to early broadband (0-100 Hz) phase and power RDMs, followed by sustained gamma power (>60Hz), and beta power (20-30Hz) at stimulus offset. The second cluster corresponded to broadband (0-100Hz) and sustained (0.1-0.4s) phase effects after stimulus onset, without any noticeable power effects. The third cluster consisted primarily of sustained (0.1-0.6s) beta (10-30Hz) and low-gamma (<60Hz) power, without any noticeable phase effects. The fourth cluster reflected broadband phase effects (0-100Hz) at stimulus offset (without associated power effects). The last 3 clusters (5-7) all displayed pre-stimulus effects in alpha-beta power, or alpha or gamma phase, characteristic of spontaneous, stimulus-unrelated activity that we did not investigate further (Figure 6). The clustering analysis performed on the MEG RDMs thus identified four main clusters of power and phase oscillatory components that occurred at different time points and in different frequency bands after stimulus onset.

**Figure 5:**
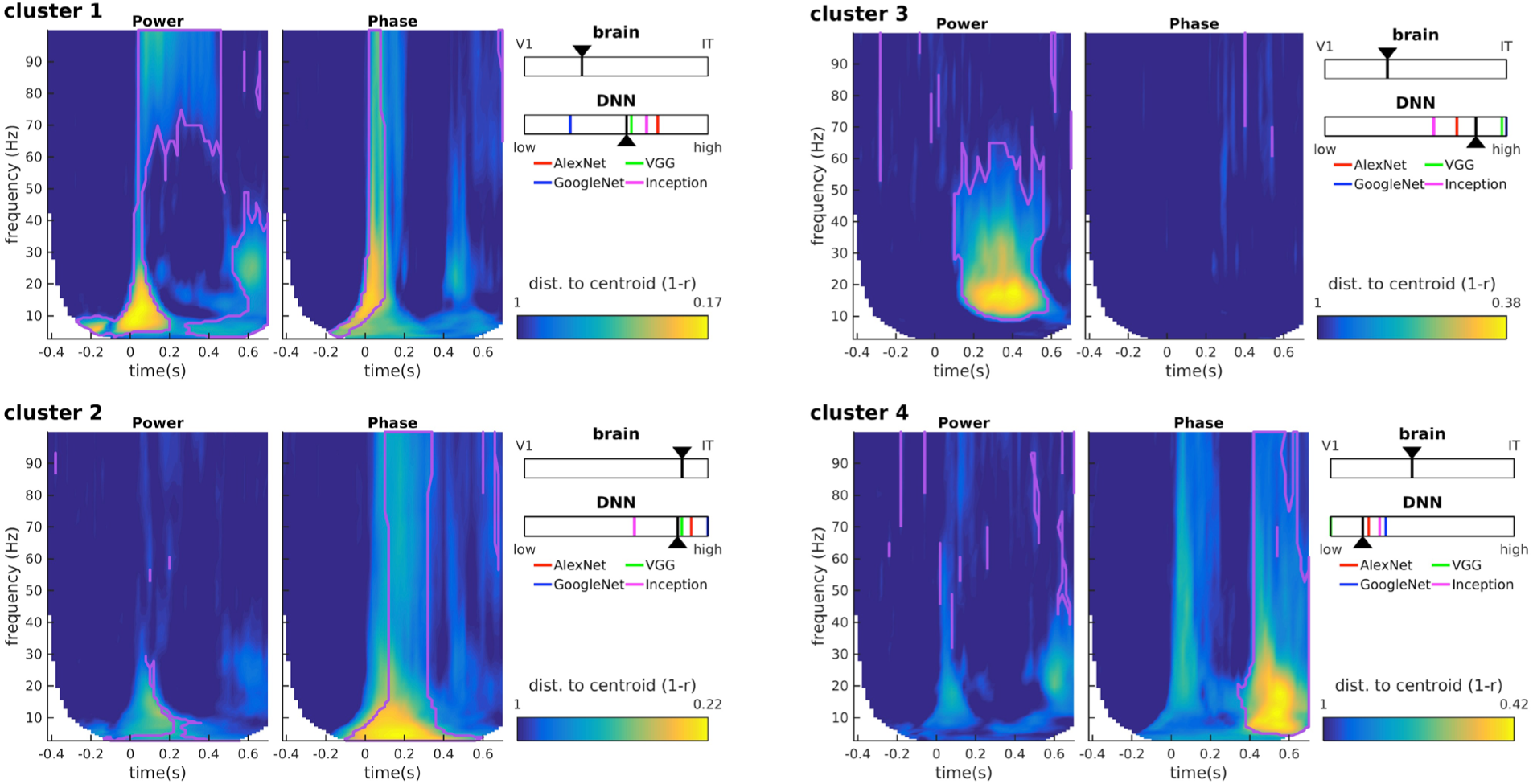
Clustering analysis. K-means clustering was performed on the MEG power and phase RDMs. Each time-frequency plot shows the distance of each RDM from the centroid of the cluster. The purple lines correspond to the cluster boundaries as returned by the k-means algorithm, indicating that all points within the purple lines are assigned to this specific cluster based on their distance to the different cluster centroids. The insets show the relative degree of RSA between the cluster centroid and V1/IT (top), or the cluster centroid and the DNN layer hierarchy (bottom). For the DNNs, the layer with maximum RSA, normalized by the number of layers in the DNN hierarchy, and averaged across the four DNN types (colored ticks), was taken as the layer that corresponded to each cluster centroid (black arrowhead).

**Figure 6:**
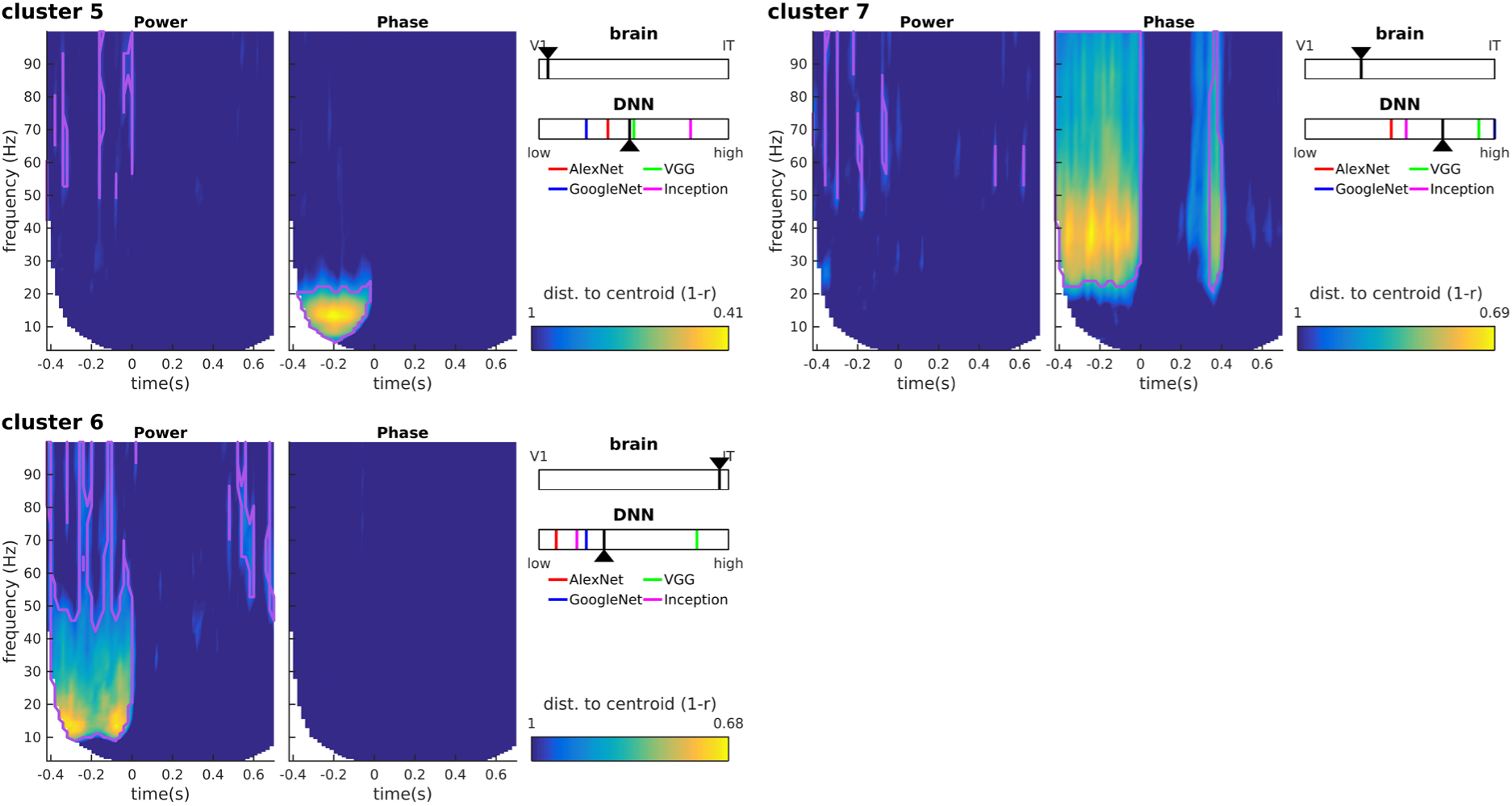
Clustering results for clusters 5-7. We identify these clusters as noise components because (i) their distance to the cluster centroid is typically higher than for other clusters, and (ii) they mainly map onto pre-stimulus oscillatory activity. Pre-stimulus oscillations, while accounting for a sizeable portion of the (notoriously noisy) MEG signal variance, cannot possibly encode the identity of a stimulus that has not been presented yet. Pre-stimulus alpha is a well-studied oscillatory component reflecting the attention state of the observer, and whose phase is known to modulate the subsequent ERP amplitudes and latencies; as such, it is not surprising that the phase of this oscillatory component would induce a separate cluster of RDM patterns (cluster 5). Similarly, pre-stimulus alpha-beta power (cluster 6) and gamma phase (cluster 7) could reflect preparatory attention or motor signals (including muscular artifacts) not related to stimulus identity. Notations as in Figure 5.

How do the oscillatory representations in each cluster, and their different time and frequency profiles relate to local processing in V1 and IT as measured by fMRI representations? To address this question, we performed RSA between the cluster centroids and the V1 and IT RDMs. The cluster centroids correlated to different degrees with both V1 and IT (all partial *R*-values between 0.12 and 0.49; all significant at *p* < 1e-5 with a surrogate test; see methods). To evaluate the relative importance of each area’s representational similarity with the cluster centroid, we combined the two RSA partial *R*-values into a single scale (see Methods, equation 1). According to this scaling (see insets in Figure 5), the transient broadband phase and power effect with sustained gamma power in cluster 1 corresponded best with V1 representations (i.e., “low-level”). Conversely, the broadband sustained phase effects of cluster 2 corresponded best to IT representations (“high-level”). The other two clusters (sustained beta-gamma power in cluster 3, broadband offset-transient phase in cluster 4) had more balanced similarity to both V1 and IT, with a slight inclination towards V1. Thus, transient oscillatory components occurring around stimulus onset correspond more closely to V1 representations, whereas the more sustained components could be either more IT-like, or less localized depending on the frequency of the oscillations. These results thus suggest a complex link between oscillatory representations and local processing in V1 or IT. To try to clarify these relationships we next turned to using deep neural networks as a template for object representations.

### Assessing representational complexity with deep neural networks

The fMRI RDMs are a representation of the image set in the multi-voxel space of V1 and IT. However, these fMRI representations are static because the fMRI BOLD signal used to construct the RDMs was measured over a period of several seconds. Neuronal activity in these regions, on the other hand, is known to evolve over fairly rapid timescales, on the order of hundreds of milliseconds as a result of feedback and top-down signals (Roelfsema, Lamme et al. 1998, Lamme and Roelfsema 2000). The fMRI RDMs are thus limited representations of the image set, potentially mixing basic and refined brain activity from different moments in each trial. Therefore, while it is tempting to interpret the oscillatory signals composing cluster 1 as basic in complexity, because they are more V1-like, and those forming cluster 2 as refined (more IT-like), such a conclusion would be premature as it ignores the dynamics of neural responses within and across brain regions, and how these neural responses evolve over different timescales. To obtain a complementary picture of low and high-level object representations, we considered the representations of our image set in different layers of feed-forward deep neural networks (DNNs) pretrained on a large dataset of natural images. To ensure the generality of our results we assessed four different DNNs: (AlexNet (Krizhevksy, Sutskever et al. 2012), VGG16 (Simonyan and Zisserman 2014), GoogleNet (Szegedy, Liu et al. 2015), and InceptionV3 (Szegedy, Ioffe et al. 2017)). Activity in each layer of these DNNs is not influenced by top-down or recurrent connections, and consequently represents a truly hierarchical evolution in the complexity of image representations, from basic to refined. Indeed, several studies have suggested that DNN representations approximate the feed-forward cascade of the visual processing hierarchy in the brain (Khaligh-Razavi and Kriegeskorte 2014, Cichy, Khosla et al. 2016). Performing RSA between MEG oscillatory RDMs and DNN layer RDMs should thus reveal which features of the MEG oscillatory representations correspond to basic vs. refined object representations.

An RDM was obtained for each convolutional layer of the four DNNs. RSA was then performed between the cluster centroids of the MEG RDMs and the DNN RDMs. For each cluster and DNN, the layer with maximum RSA was determined, and scaled between 0 (lowest layer, basic information) and 1 (highest layer, refined information) based on the number of layers in the DNN hierarchy. Despite slight differences between the four DNNs, the analysis revealed that clusters 2 and 3 mapped best to higher DNN layers, cluster 1 to intermediate layers, and only cluster 4 had similarity to lower layers. This is in stark contrast with the results of fMRI RSA, which had ranked clusters 2, 4, 3 and 1 in order of decreasing complexity. The most striking difference is obtained for cluster 3 (sustained beta-gamma power): a high-level refined representation according to DNNs, but closer to V1 than to IT according to fMRI. Based on the logic above, this cluster is likely to reflect feed-back signals that carry refined object information (visible in high DNN layers) down to lower brain regions (visible in V1 BOLD signals).

## Discussion

Our results show that MEG oscillatory components at different frequencies carry stimulus-related information at specific times, which can be linked, via RSA, to stimulus representations in different brain regions (V1, IT), and with different representational complexity (as measured by deep neural networks). Importantly, the representational dynamics of brain oscillations can be very differently expressed by power vs. phase signals. At stimulus onset and offset, broadband phase transients (possibly related to fluctuations in evoked potential latencies) carry mainly basic or intermediate-complexity information (clusters 1, 4 in Figure 5). However, during stimulus presentation, sustained phase information is visible across all frequencies, and consistently maps to high-level refined representations (IT and high DNN layers, cluster 2). Oscillatory power components (clusters 1 and 3) tend to correlate with both V1 and IT fMRI representations (with an inclination towards V1); however, onset-transient low-frequency (<20Hz) power together with sustained high-frequency (>60Hz) power (i.e., cluster 1) correspond best to intermediate DNN layers, whereas sustained beta-gamma power (20-60Hz) clearly maps to the highest DNN layers (cluster 3).

In our study we found no simple mapping between low/high-level (or basic/refined) representations and oscillatory components (power/phase) or frequency. Both low-frequency (theta, alpha) and high-frequency (beta, gamma) oscillatory signals can carry either low/basic or high-level/refined representations at different times (e.g. clusters 2 vs. 4). Similarly, both phase and power signals can carry either low or high-level representations (e.g. clusters 1 vs. 3). The picture that emerges is a rather complex one, in which successive interactions between different oscillatory components in different brain regions and at different frequencies reflect the different stages of neural processing involved in object recognition.

Our results highlight the importance of complementing MEG-fMRI RSA with another measure of representational content such as feed-forward DNNs (Hebart, Bankson et al. 2018, Khaligh-Razavi, Cichy et al. 2018). fMRI BOLD signals are often analyzed such that they reflect a single static representation. Thus, they cannot distinguish dynamics in local patterns as for example early feedforward and later feedback activity. By design, feedforward DNN layers cannot be dynamically influenced by feedback signals, and could be considered to provide a template for basic vs. refined representations during the different stages of image processing. Perhaps the best illustration of this notion stems from the discrepancy between fMRI and DNN RSA for MEG cluster 3, which suggests that sustained beta-gamma power during stimulus presentation could reflect feedback signals: best corresponding to V1 fMRI activity (low-level), but higher DNN layers (refined). Without this additional information (e.g., looking at Figure 3a alone), one might have interpreted sustained beta-power as a strictly low-level signal. The observed distinction between sustained power effects at lower frequencies (beta and low-gamma, cluster 3) vs. higher frequencies (high-gamma, cluster 1) is consistent with a large number of recent studies that reported a functional distinction between gamma-band and beta-band signals, respectively supporting feed-forward and feedback communication (Fontolan, Morillon et al. 2014, van Kerkoerle, Self et al. 2014, Bastos, Vezoli et al. 2015, Michalareas, Vezoli et al. 2016). Future work could attempt to separate feedforward from feedback signals (e.g. with backward masking), to confirm the differential contribution of gamma and beta-band oscillatory frequencies to feedforward vs. feedback object representations, as determined with RSA.

In addition to their involvement in the transmission of feedforward and feedback signals, several studies have shown that different oscillations can carry distinct information about stimulus properties (Hebart, Bankson et al. 2018, Khaligh-Razavi, Cichy et al. 2018). Here we considered whether oscillatory components in different frequencies correspond to basic or more refined stimulus processing stages. Our results suggest that most oscillatory brain activity, at least at the broad spatial scale that is measured with MEG, reflects already advanced stimulus processing in object detection tasks. This result can be seen in Figure 5 where most oscillatory components are more related to higher-level DNN layer representations, with the exception of the offset-transient (cluster 4). Indeed, one might have expected that stimulus representations at both stimulus onset and offset are more reflective of transient low-level/basic processing. However, while both onset and offset signals (cluster 1 and cluster 4) are better matched to V1 than IT (“low-level”, see also Figure 3c, d), in terms of DNN activations the offset-transient (cluster 4) appears to be much more basic in complexity and the onset-transient more refined (cluster 1). A tentative explanation could be that the continued presence of the stimulus after the onset-transient supports a rapid refinement of object representations, which would not be the case for the offset-transient (because the stimulus is absent from the retina). Indeed, it is remarkable that, aside from this offset-transient broadband phase activity (cluster 4), no other oscillatory signal was found to reflect low-level DNN layers (i.e., basic information).

In conclusion, our results help characterize the representational content of oscillatory signals during visual object perception. By separately considering hierarchical level (V1/IT) and representational complexity (based on DNNs), we narrow the gap between whole-brain oscillations and visual object representations supported by local neural activation patterns.

## Notes

### Competing Interest Statement

The authors have declared no competing interest.

